# Mass Mortality of Marine Mammals Associated to Highly Pathogenic Influenza Virus (H5N1) in South America

**DOI:** 10.1101/2023.02.08.527769

**Authors:** Víctor Gamarra-Toledo, Pablo I. Plaza, Roberto Gutiérrez, Giancarlo Inga-Diaz, Patricia Saravia-Guevara, Oliver Pereyra-Meza, Elver Coronado-Flores, Antonio Calderón-Cerrón, Gonzalo Quiroz-Jiménez, Paola Martinez, Deyvis Huamán-Mendoza, José C. Nieto-Navarrete, Sandra Ventura, Sergio A. Lambertucci

## Abstract

We report a massive mortality of more than 3,000 sea lions (*Otaria flavescens*) of Peru associated with a Highly Pathogenic Influenza Virus (H5N1). The transmission pathway of H5N1 may have been through the close contact of sea lions with infected wild birds. We cannot rule out direct transmission among sea lions.

---

The recent panzootic event (2020-2022) caused by the highly pathogenic avian influenza (HPAI) A (H5N1) is the largest observed so far, several global outbreaks having been caused (1). At the end of 2022, the H5N1 virus reached South America (Peru, Ecuador, Colombia, Venezuela and Chile), with alarming bird mortalities in Peru (2). This pathogen was detected for the first time in wild birds in Peru on 13 November (2). Reports suggest the virus killed more than 50,000 wild birds by the end of 2022, particularly Peruvian pelicans (*Pelecanus thagus*) and Peruvian boobies (*Sula variegata*) (2,3). The large biomass of infected wild birds may have led to a spillover event affecting predators and scavengers, including marine mammals cohabiting with them, as reported in other parts of the world (4). Here, we report the death of 3,108 sea lions (*Otaria flavescens*) on Peruvian coasts over 5 weeks and necropsies of some individuals that suggest they were affected by the Highly Pathogenic Avian Influenza Virus (H5N1), this was lately confirmed by government reports.

During January and February 2023, we performed detailed survey of died mammals in marine protected areas of Peru with SERNANP personal (Servicio Nacional de Áreas Naturales Protegidas por el Estado), throughout a surveillance system for avian influenza in high biodiversity areas. More than three thousand sea lions were found dead or dying on Peruvian beaches (Fig. 1.1, Table 1). The high mortality observed was worrisome; for instance, up to 100 dead individuals were found floating together in the sea, or 1,112 individuals died just in one island (Isla San Gallan); one of the places with the highest populations of sea lions in Peru. Those are unprecedented observations for the Southern Hemisphere (Fig. 1.1, Fig. 1.2 A and B, Table 1). The clinical signs of dying individuals were mainly neurological, such as tremors, convulsions and paralysis. They also showed respiratory signs such as dyspnea, tachypnea, nasal and buccal secretions and pulmonary edema (Fig. 1.2 C). At the time of the deaths many sea lions were pregnant or recently calved females; several abortions were observed. There are no other records of such high mortality of aggregated sea lions, and this is the highest mammal mortality recorded in the world that can be associated to HPAV until now.

**Figure 1.**
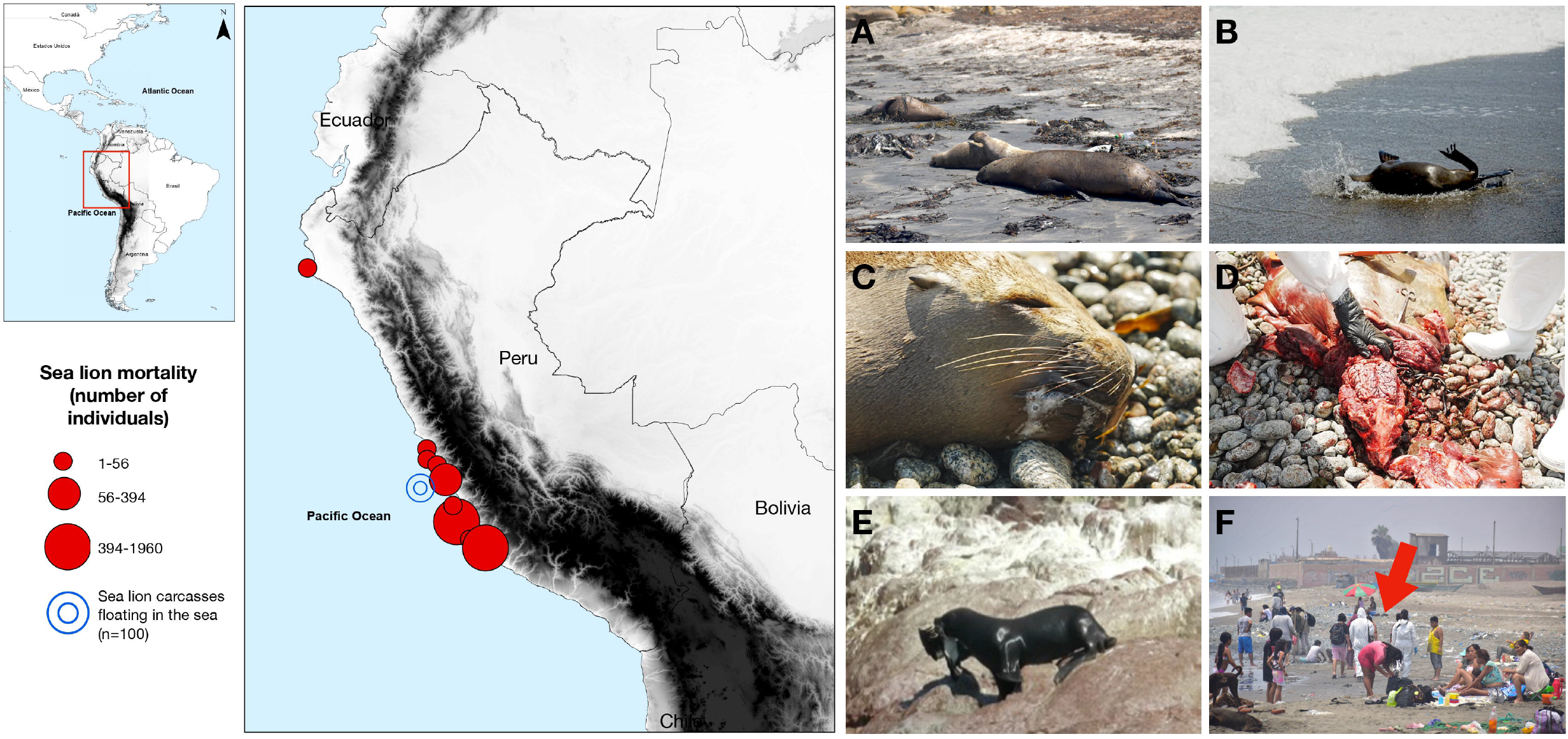
**1.** Map of geographical distribution of high mortalities in sea lions in January and February 2023 on the Peruvian coastline. **2.** Images showing the field work in the Paracas National Reserve on the Peruvian coastline, February 2023. A) Sea lion carcasses on the beach; B) Dying sea lion with ataxia; C) Dead sea lion with avian flu symptoms (whitish secretions); D) Sea lion necropsy showing a congestive brain; E) Sea lion trapping and eating an infected Guanay cormorant on January 23th of 2023 in the Reserva Nacional Paracas; F) Field work sampling on a beach with a large number of bathers in the surroundings of infected carcasses (the red arrow indicates SERNANP personal with health protection equipment conducting field survey). Photo credits: A, B and D, Daniel Ampuero; C and F, Giancarlo Inga; E, Sandra Lizarme.

**Table 1.**
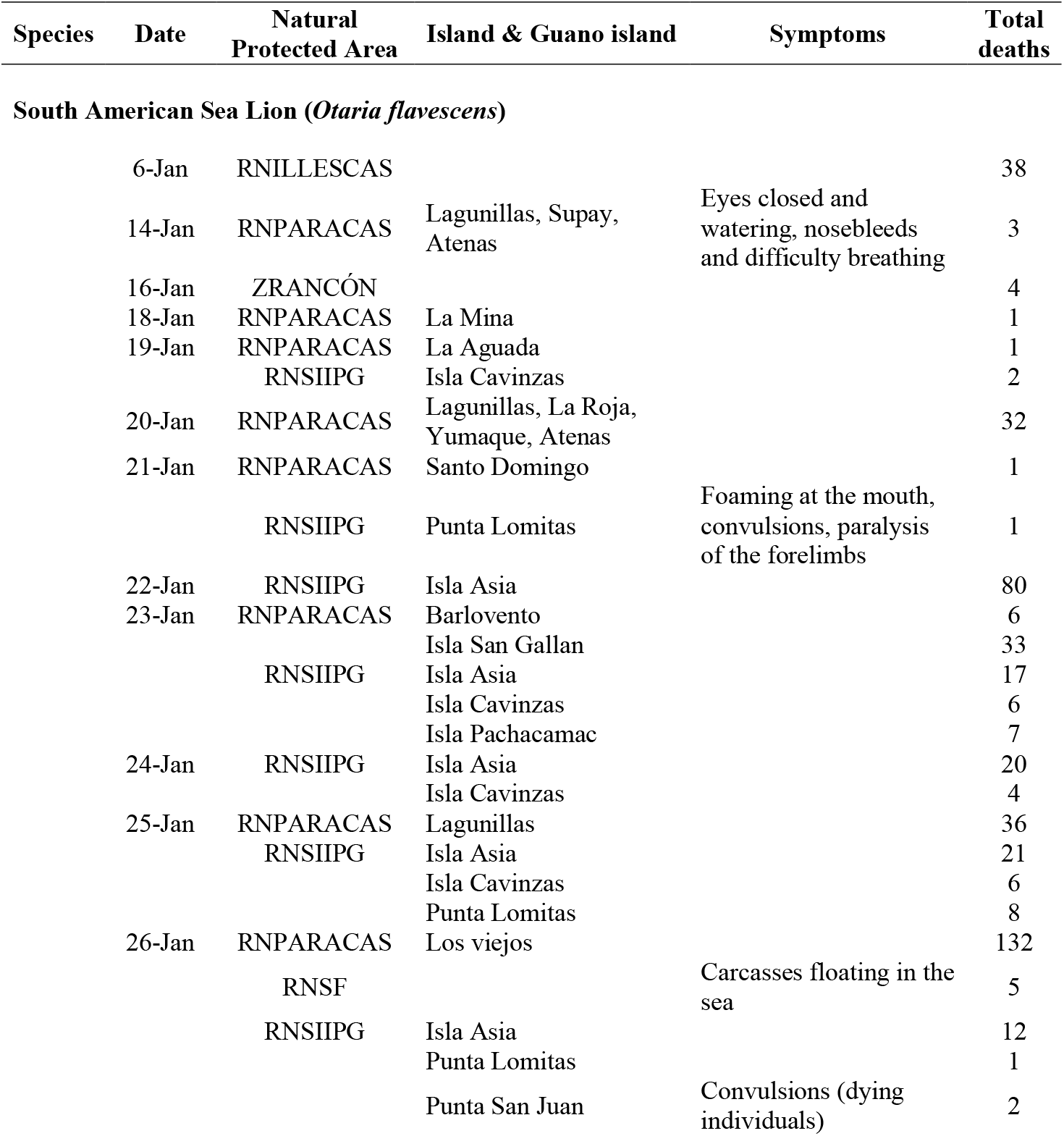
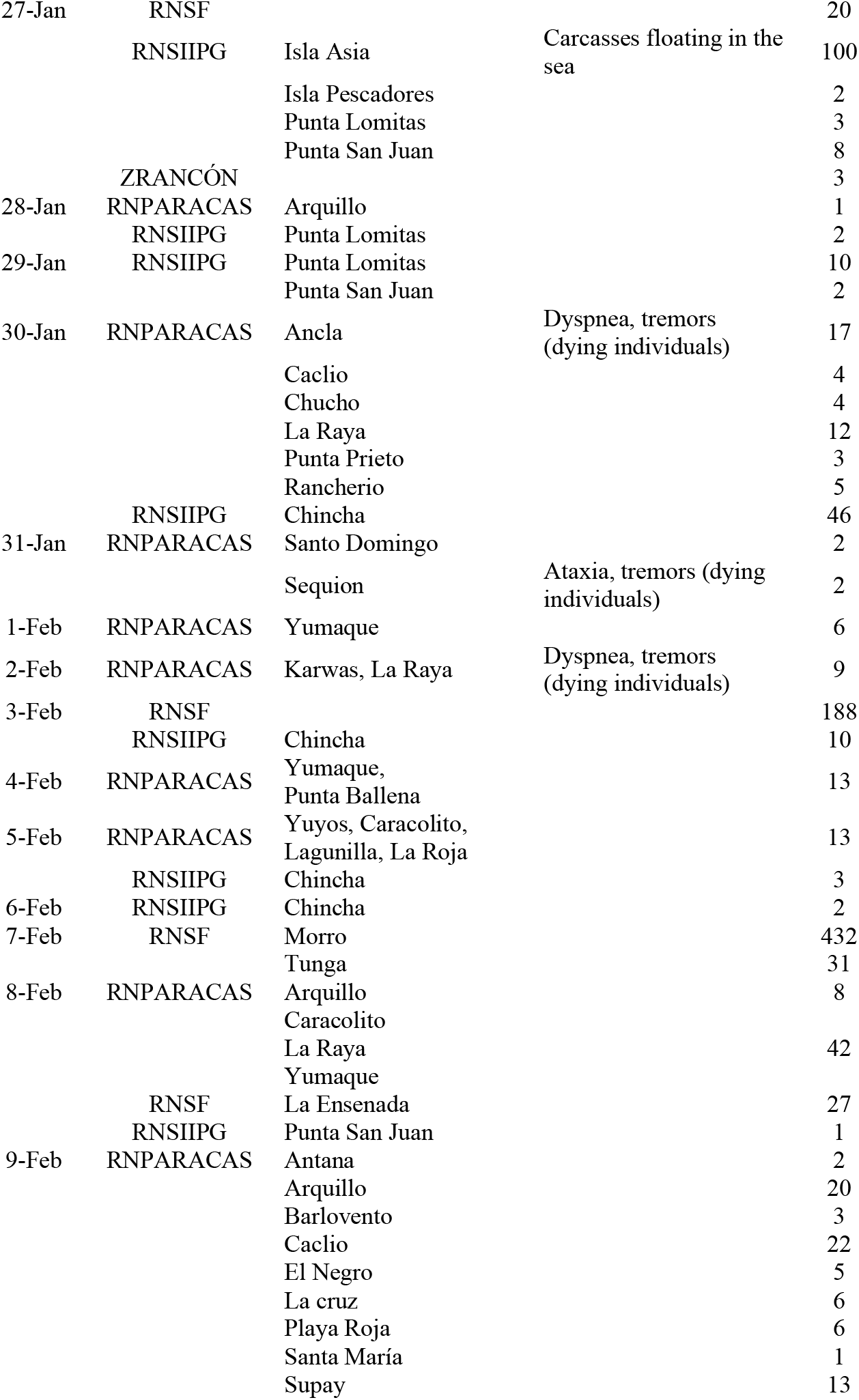
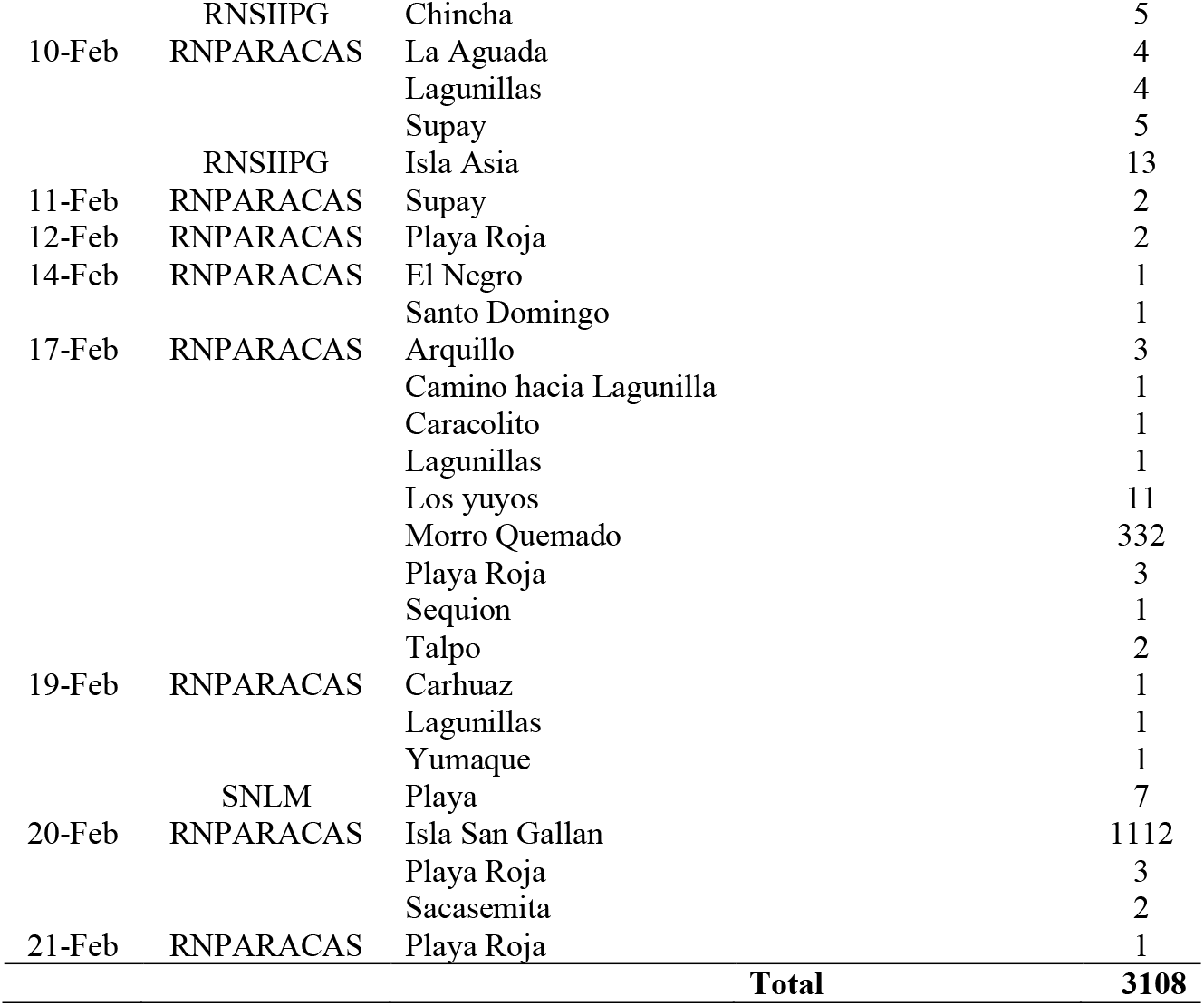
Sea lion mortality associated with Highly Pathogenic Avian Influenza Virus (H5N1) in protected areas of Peru between January and February 2023. Abbreviations for the Natural Protected Areas are: RNSF = Reserva Nacional San Fernando, RNILLESCAS = Reserva Nacional Illescas, RNPARACAS = Reserva Nacional Paracas, ZRANCÓN = Zona Reservada Ancón, RNSIIPG = Reserva Nacional Sistema de Islas, Islotes y Puntas Guaneras.

Individuals were examined by veterinarians and some dead animals were necropsied. The body condition of the sea lions necropsied ranged from good to very good, suggesting they died due to an acute health problem. Substantial quantities of whitish secretions filling the upper respiratory tracts (trachea and pharynx) were observed in the necropsies and in dying animals (Fig. 1.2 C), which explains the severe dyspnea and tachypnea clinically observed by veterinarians. Lungs were congestive, with hemorrhagic focus compatible with interstitial pneumonia. Brains were also congestive with hemorrhagic focus compatible with encephalitis, which explains the neurological signs observed in dying individuals (Fig. 1.2 D); this agrees with the result of other study on mammals infected with H5N1 (5). The small intestine showed necrotic focus compatible with duodenitis. At the time of writing this letter (February 2023) sea lions mortalities continue and have reached up to 3,108 died individuals. Given the epidemiologic situation produced by H5N1 in wild birds that cohabit with these sea lions (2,3), our most plausible diagnosis was acute disease caused by HPAV. Official information from Peruvian government confirmed that some of those sea lions tested positive to HPAI (H5N1) (3,6,7).

In conclusion, sea lions of Peru developed a deadly associated disease that have produced massive mortality in several regions of the Peruvian coastline (Fig. 1.1). The sea lion mass mortality described is compatible with systemic HPAI that resulted in acute encephalitis and pneumonia. Some specific cases of infection and mortality of marine mammals due to HPAV with similar clinical and anatomopathological characteristics have been reported in previous studies around the world (4,8,9).

The source of the HPAI affecting these sea lions was most probably the large number of infected alive birds or their carcasses on the Peruvian coastline (2,3). Sea lions may be infected by close contact with these carcasses and even through their consumption (see an example of a sea lion feeding on a sick bird in Fig. 1.2 E). However, based on recent research suggesting the first mammal-to-mammal infection in *Neovison vison* (10) and the large number of sea lions currently affected at the same time, we cannot rule out that the virus has adapted to mammals and that sea lion-sea lion transmission has begun in Peru, and this needs to be urgently studied.

Worryingly, at the time of this submission Bolivia, Uruguay and Argentina add to list of countries that have reported HPAV in their territories for the first time (11). Moreover, authorities from Chile have reported the first sea lion dead due to HPAV in the north of the country with similar clinical signs that we reported in Peru (11).

Further research is required to address the transmission pathway in this social species. We would like to call attention to the fact that in this geographical region of the world, human–infected animal interaction is common (e.g., people stain in beaches with sea lions carcasses, (12) (Fig. 1.2 F), so infections might begin to rise and this must be addressed if we are to avoid the risk of a pandemic.

## Acknowledgments

We thank the Unidad Operativa Funcional Monitoreo, Vigilancia y Control (Dirección de Gestión de Áreas Naturales Protegidas) of the Servicio Nacional de Áreas Naturales Protegidas por el Estado (SERNANP) of the Peruvian Ministry of Environment, for the permission to access and use the information provided (Expediente Tupa N° 0108-2023; CARTA No 053 - 2023-SERNANP-AIP). We especially thank SERNANP park rangers for their help in the field survey and their involvement in the conservation and monitoring of wildlife in response to this health emergency. We also thank the staff of the Wildlife Conservation Society (WCS), especially Paulo Colchao, Jorge Martinez and Mariana Montoya, for their institutional and logistical support during field sample collection. Finally, we thank all public institutions that contribute to the monitoring of this epizootic and epidemiological surveillance in Peru.

## Biography

Víctor Gamarra-Toledo, MSc(c). Biologist, research associate at the Museum of Natural History (MUSA), Universidad Nacional de San Agustín de Arequipa, Peru. With experience in wildlife conservation in coastal ecosystems of Peru.

